# Single-molecule FRET monitors CLC transporter conformation and subunit independence

**DOI:** 10.1101/2020.09.07.286831

**Authors:** Ricky C. Cheng, Ayush Krishnamoorti, Vladimir Berka, Ryan J Durham, Vasanthi Jayaraman, Merritt Maduke

## Abstract

“CLC” transporters catalyze the exchange of chloride ions for protons across cellular membranes. As secondary active transporters, CLCs must alternately allow ion access to and from the extracellular and intracellular sides of the membrane, adopting outward-facing and inward-facing conformational states. Here, we use single-molecule Förster resonance energy transfer (smFRET) to monitor the conformational state of CLC-ec1, an *E. coli* homolog for which high-resolution structures of occluded and outward-facing states are known. Since each subunit within the CLC homodimer contains its own transport pathways for chloride and protons, we developed a labeling strategy to follow conformational change within a subunit, without crosstalk from the second subunit of the dimer. Using this strategy, we evaluated smFRET efficiencies for labels positioned on the extracellular side of the protein, to monitor the status of the outer permeation pathway. When [H^+^] is increased to enrich the outward-facing state, the smFRET efficiencies for this pair decrease. In a triple-mutant CLC-ec1 that mimics the protonated state of the protein and is known to favor the outward-facing conformation, the lower smFRET efficiency is observed at both low and high [H^+^]. These results confirm that the smFRET assay is following the transition to the outward-facing state and demonstrate the feasibility of using smFRET to monitor the relatively small (~1 Å) motions involved in CLC transporter conformational change. Using the smFRET assay, we show that the conformation of the partner subunit does not influence the conformation of the subunit being monitored by smFRET, thus providing evidence for the independence of the two subunits in the transport process.

**SUMMARY:** Cheng, Krishnamoorti et al. use single-molecule Förster energy resonance transfer measurements to monitor the conformation of a CLC transporter and to show that the conformational state is not influenced by the neighboring subunit.

## INTRODUCTION

Members of the CLC (“Chloride Channel”) family are found at all levels of biological complexity, from single-cell bacteria to humans (Jentsch and Pusch, 2018). The family includes both passive ion channels and active transporters. The transporter homologs exchange chloride (Cl^−^) for protons (H^+^). These homologs reside in intracellular membranes, including lysosomes, endosomes, osteoclasts, and synaptic vesicles, where they play critical roles in regulating [Cl^−^] and [H^+^], and where mutations lead to severe pathologies (Poroca et al., 2017; Jentsch and Pusch, 2018; Nicoli et al., 2019; Gianesello et al., 2020). The CLC channel homologs are expressed in the plasma membranes of essentially all cells, carrying out functions ranging from voltage-dependent signaling to epithelial ion transport (Fahlke and Fischer, 2010; Denton et al., 2013; Wang et al., 2017; Fernandes-Rosa et al., 2018; Jentsch and Pusch, 2018; Elorza-Vidal et al., 2019; Stowasser et al., 2019; Altamura et al., 2020).

All CLCs are homodimeric, with each subunit having its own pathways for the substrate ions (Cl^−^ and H^+^) (Accardi, 2015; Jentsch and Pusch, 2018). The Cl^−^ pathway is formed by the convergence of helix dipoles and a conserved intracellular loop, and it is capped by a key glutamate residue, termed “Glu_ex_” (**Figure 1A**). Glu_ex_ acts as both a “gate” to the Cl^−^-permeation pathway and as a H^+^-transfer residue. The passage of Cl^−^ and H^+^ occurs along pathways that are shared at the extracellular portion of the protein and then diverge towards the intracellular side (Accardi et al., 2005; Accardi, 2015; Chavan et al., 2020). In the channel homologs, H^+^ transfer is uncoupled from Cl^−^ permeation, such that thousands of Cl^−^ ions are transported for every H^+^, and thus H^+^ transport can be measured only indirectly (Lisal and Maduke, 2008, 2009). In the transporters, stochiometric (2:1) Cl^−^/H^+^ coupling is observed under a wide range of conditions (Accardi et al., 2004; Accardi and Miller, 2004; Scheel et al., 2005; Accardi, 2015).

**Figure 1.**
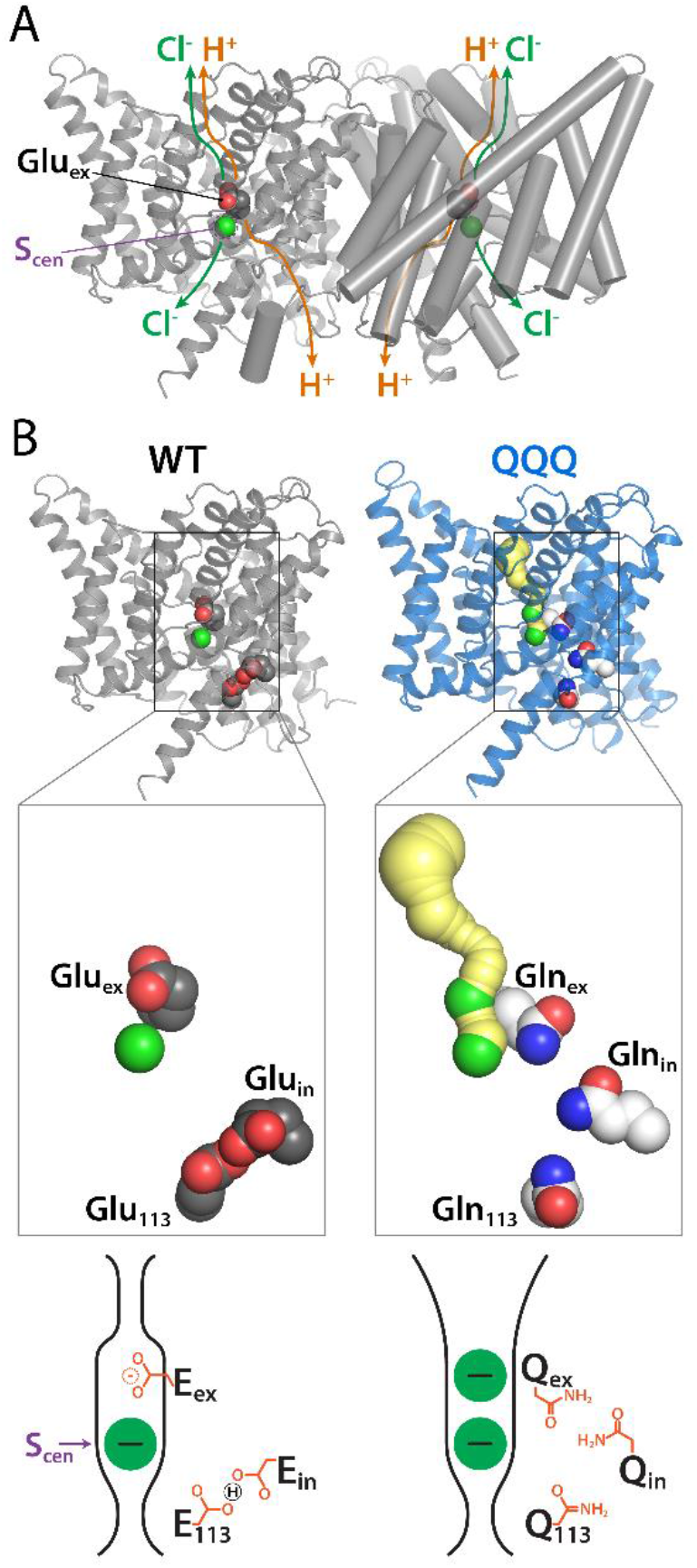
CLC structure overview. (**A**) Structure of CLC-ec1, illustrating the homodimeric architecture and the Cl^−^ and H^+^ permeation pathways (one per subunit). Glu_ex_, shown in spacefill, is a key residue in both CLC channels and transporters. In the channels, it acts as a gate to the Cl^−^-permeation pathway; in the transporters, it acts as both a gate to the Cl^−^-permeation pathway and as a H^+^-transfer residue. Glu_ex_ directly blocks Cl^−^ in the central site (S_cen_) from the extracellular solution. (**B**) Comparison of WT CLC-ec1 (PDB 1OTS, occluded state) to the “QQQ” CLC-ec1 mutant (PDB 6V2J, outward-facing state). The three glutamate residues that are mutated to glutamine in the QQQ protein are shown spacefilled in each structure. Cl^−^ pathways, shown in yellow, were detected using Caver (Chovancova et al., 2012) with probe radius 1.2 Å, starting from the Cl^−^ ion at the S_cen_ site (lower of the two Cl^−^ sites shown in the QQQ structure). The QQQ mutant has a pathway from S_cen_ to the extracellular side, while the WT protein does not. Expanded views below highlight the Cl^−^ pathway and the changes in positioning of the three glutamate/glutamine residues. In QQQ, Gln_ex_ is rotated to a position that can accept H^+^ from the intracellular side, with Gln_in_ and Gln_113_ separated from one another, providing space for water/protons to flow to and from the intracellular side. The bottom panel shows cartoon depictions of the occluded and outward-facing states.

CLC-ec1, a homolog from *E. coli*, provided the first high-resolution CLC structure and has subsequently been intensely studied (Mindell et al., 2001; Dutzler et al., 2002; Dutzler, 2007; Matulef and Maduke, 2007; Accardi, 2015). It is an excellent subject for investigations of structure and function, as it is highly amenable to biochemical manipulations and is well validated as a model for understanding mammalian homologs (Matulef and Maduke, 2007; Accardi, 2015; Jentsch and Pusch, 2018). Several high-resolution structures of CLC-ec1, under varying experimental conditions and with varying mutations, are available. Most of these show the same overall conformation – an “occluded” state in which the bound Cl^−^ substrate is blocked from both the extracellular and intracellular solutions. Recently, a crystal structure of a CLC-ec1 triple mutant (“QQQ” – to mimic protonation of three key glutamate residues) revealed the first molecular details of the CLC-ec1 outward-facing conformational state, with the permeation pathway open for Cl^−^ to exchange to the extracellular side (Chovancova et al., 2012; Chavan et al., 2020) (**Figure 1B**). In addition, the structure involves a rearrangement of the three glutamine residues: Glu_ex_ (Gln_ex_), the H^+^-transfer residue, reaches inwards for H^+^ exchange to the intracellular side, while the intracellular Gln residues separate from one another, allowing for a water/H^+^ pathway from the intracellular side (**Figure 1B**). The relevance of this conformation to the wild-type (WT) protein was confirmed using double electron-electron resonance (DEER) spectroscopy experiments to compare conformations of WT and QQQ mutant proteins; these experiments revealed that wild-type CLC-ec1 adopts a QQQ-like conformation when the pH is lowered from 7.5 to 4.5 to protonate the key glutamate residues (Chavan et al., 2020).

Spectroscopic approaches are powerful tools for monitoring protein conformational dynamics that are not captured in static structures (Elvington and Maduke, 2008; McHaourab et al., 2011; Sekhar and Kay, 2019; Sahu and Lorigan, 2020). Single-molecule Förster resonance energy transfer (smFRET) has emerged as an important player in the spectroscopic toolbox, as a method that can monitor conformational transitions in real time (Zhao et al., 2010; Landes et al., 2011; Akyuz et al., 2015; Dolino et al., 2015; Vafabakhsh et al., 2015; Juette et al., 2016; Dyla et al., 2017; Han et al., 2017; Lerner et al., 2018; Ren et al., 2018; Wang et al., 2018; de Boer et al., 2019; MacLean et al., 2019; Mazal and Haran, 2019; Zhu et al., 2019; Carrillo et al., 2020; Durham et al., 2020). In addition, the single-molecule approach confers flexibility in labeling strategies, allowing labeling at multiple sites without concerns of sample heterogeneity, which can thwart ensemble-spectroscopic measurements. For example, the DEER measurements in our previous study used a labeling strategy with one label per subunit, measuring inter-subunit distance distributions (Chavan et al., 2020); this strategy is valuable because it avoids spectral overlap of intra- and inter-subunit distance distributions that would occur with multiple labels per subunit, but it does not provide information about independent movements within a subunit. Here, we design smFRET experiments to monitor conformational change within one subunit of the CLC-ec1 dimer, without interference from labels on the second subunit. Using this strategy, we demonstrate that the conformation at the CLC-ec1 extracellular gate can be monitored as a function of pH and that this conformational change is independent of the conformation adopted by the second subunit.

## MATERIALS AND METHODS

### Protein Overexpression and Purification

Mutagenesis experiments to generate the E377C/H234C, Y75C/H234C, and Y75C/E377C mutants were carried out in a cysteine-less background (C85A/C302A/C347S) (Nguitragool and Miller, 2007), in either an otherwise WT CLC-ec1 background or in a “QQQ” background (E148Q/E203Q/E113Q) (Chavan et al., 2020). Mutants were verified by sequencing the entire coding region of the gene. Strep-tagged CLC-ec1 was generated using Gibson assembly protocol (Gibson et al., 2009). Transformation, expression, and purification of CLC-ec1 was performed as previously described (Accardi et al., 2004), but with the following changes. For the expression of CLC-ec1 heterodimers, two constructs were used in plasmid transformation at equal proportion. To ensure that only cells with both constructs survive, the double-cysteine mutant constructs contained an ampicillin resistance gene, and the cysteine-less/strep-tagged construct contained a kanamycin-resistance gene. Cells from the transformation plate were used to inoculate 1 L of Terrific broth (Sambrook et al., 1989) in a 4-L unbaffled flask with 100 μg/L of ampicillin and 50 μg/L of kanamycin. Protein expression was induced with 0.2 mg/L of anhydrotetracycline when the OD_550_ reached between 1.6 and 2.0. Dithiothreitol (DTT) was added to the media at the time of induction (0.8 g/L) and ~ 1.5 h after induction (0.5 g/L). During purification, all buffers used during intermediate steps contained 1 mM DTT, and the buffer for the final size-exclusion chromatography step (10 mM Na-HEPES, 150 mM NaCl, 5 mM decyl maltoside, pH 7.5) contained 1 mM Tris(2-carboxyethyl)phosphine hydrochloride (TCEP) to preserve the free thiols. The purified protein was shipped by FedEx (Standard Overnight) with ice packs in a Styrofoam box to the Jayaraman lab for fluorophore-labeling and smFRET measurements.

### Functional Assays

Purified CLC-ec1 samples were reconstituted into liposomes according to previous protocols (Walden et al, 2007; Chavan et a l, 2020). *E. coli* polar lipids (Avanti Polar Lipids, Alabaster, AL) (in chloroform) in a round-bottomed flask were dried under argon gas. To further remove chloroform, the dried lipids were then dissolved in pentane and dried again under vacuum on a rotator, followed by further drying under argon for 5 minutes. The lipids were then resuspended in buffer R1 (333 mM KCl, 55 mM Na-citrate, 55 mM Na_2_HPO_4_, pH 6.0) with 35 mM CHAPS at 20 mg lipids per mL, with rotation for 1.5 to 2 hours. Purified protein samples (at 0.4 μg protein per mg lipid) or control solutions (containing no protein) were added to the prepared lipid-detergent mix and incubated for 10-20 min. Each protein or control reconstitution was dialyzed in buffer R1 to remove detergent and allow liposome formation over 36-48 hours, with 3 buffer changes. The resulting liposomes were aliquoted for pH adjustments in duplicate. The target pH (4.0 – 7.5) was reached by further dialyses (overnight with one buffer change) in buffer R2. Each buffer R2 for a specific target pH was prepared by mixing buffer R1 with an adjustment buffer at 9:1 ratio. The adjustment buffers were prepared by varying the ratio of 1.0 M citric acid and 0.5 M Na_3_PO_4_, as outlined in **Table 1**.

**TABLE 1.** Adjustment Buffers

| <u>Target</u><br><u>pH</u> | <u>Adjustment Buffer for R2</u> |  | <u>Adjustment Buffer for F2</u> |  |
| --- | --- | --- | --- | --- |
|  | <u>Citric Acid (M)</u> | <u>Na<sub>3</sub>PO<sub>4</sub> (M)</u> | <u>Citric Acid (M)</u> | <u>Na<sub>3</sub>PO<sub>4</sub> (M)</u> |
| 4.0 | 0.51 | 0.24 | 0.49 | 0.26 |
| 4.5 | 0.44 | 0.28 | 0.42 | 0.29 |
| 5.2 | 0.35 | 0.33 | 0.33 | 0.33 |
| 6.0 | 0.26 | 0.37 | 0.26 | 0.37 |
| 7.5 | 0.16 | 0.42 | 0.16 | 0.42 |

The reconstituted liposomes were subjected to four freeze-thaw cycles followed by extrusion 15 times using an Avanti Mini Extruder fitted with a 0.4 μm-filter (GE Healthcare, Chicago, IL). For each assay, 60-μL samples of extruded liposomes were buffer-exchanged through 1.5-mL spin columns. Spin columns consisted of Sephadex G-50 Fine resin (GE Healthcare, Chicago, IL) equilibrated with pH-adjusted buffer F2 and subjected to a pre-spin at 1100 g for 25-30 s, using a clinical centrifuge. The pH-adjusted buffers F2 were prepared from mixing buffer F1 (333 mM K-isethionate, 55 mM Na-citrate, 55 mM Na_2_HPO_4_, 55 μM KCl, pH 6.0) and Adjustments Buffer (**Table 1**) at 9:1 ratio. Samples were loaded onto the spin columns, spun at 1100 g for 90 s, diluted with 600 μL of pH-adjusted buffer F2, and used immediately in flux assays. Activity was monitored by measuring the extra-liposomal [Cl^−^] using a Ag•AgCl electrode. After ~30 s recording to monitor the baseline, bulk Cl^−^ transport through CLC-ec1 was initiated by addition of 1.7 μg/mL of valinomycin (as 0.5 mg/mL stock solution in ethanol), to allow K^+^ flux to counterbalance the electrogenic Cl^−^/H^+^ flux through CLC-ec1. At the end of each assay (~ 40 s after valinomycin addition), Triton X-100 (10 μL of 10% in water) was added to release all Cl^−^ from the liposomes and allow quantification of total Cl^−^ in each assay. Calibration curves for the Ag•AgCl electrode was performed with KCl in 27-136 nmol steps. Unitary Cl^−^ turnover rates were calculated using the initial-velocity method as outlined by Walden and Coworkers (Walden et al, 2007), using 51 kDa as the molecular weight of a CLC-ec1 subunit.

### Protein labeling

Upon arrival at the Jayaraman lab, the CLC-ec1 protein was diluted 1:10 in a buffer consisting of 10 mM HEPES pH 7.5, 150 mM NaCl, 5 mM Decyl Maltoside, and 1 mM EDTA. Stock solutions of 1 mM maleimide-linked donor and acceptor fluorophores, Alexa 555 and Alexa 647 (Invitrogen), respectively, in DMSO were pre-mixed in a separate tube and then added to the diluted protein to achieve a concentration of 600 nM Alexa 555 and 2.4 μM Alexa 647 (final DMSO <2% in the labeling reaction). The sample was then rotated at room temperature for 30 minutes while being protected from ambient light to allow the fluorophores to label the CLC protein. Labeled protein was again diluted 1:4 in a buffer consisting of 10 mM HEPES pH 7.5, 150 mM NaCl, 5 mM Decyl Maltoside, and 1 mM EDTA, and the resulting labeled CLC protein was used to prepare slides for smFRET.

### Slide Preparation

Experimental methods concerning smFRET slide preparation, data collection, and data analysis are as previously described (Durham et al., 2020). In brief, glass coverslips were used to immobilize sample molecules for smFRET measurements. The coverslips were first cleaned via bath sonication in Liquinox phosphate-free detergent (Fisher Scientific) and then acetone. Further cleaning was achieved by incubating the slides in a 4.3% NH_4_OH and 4.3% H_2_O_2_ solution at 70°C followed by plasma cleaning in a Harrick Plasma PDC-32G Plasma Cleaner. After cleaning of the slide surface was completed, the slide was treated to prepare a Streptavidin-coated surface for the attachment of CLC-ec1. This process was begun via amino silanization of the slide surface with Vectabond reagent (Vector Laboratories). This step was followed by polyethylene-glycol (PEG) treatment of the slide surface with 0.25% w/w 5 kDa biotin-terminated PEG (NOF Corp., Tokyo, Japan) and 25% w/w 5 kDa mPEG succinimidyl carbonate (Laysan Bio Inc., Arab, AL). This initial PEG treatment was followed by a secondary PEG treatment with 25mM short-chain 333 Da MS(PEG)4 Methyl-PEG-NHS-Ester Reagent (Thermo Scientific). These treatments resulted in a biotin-coated surface on the slide. Next, a microfluidics chamber consisting of an input port, a sample chamber, and an output port was constructed on the slide (Litwin et al., 2019). Imaging Buffer (1mM nDodecyl-β-D-maltoside (Chem-Impex Int’l Inc., Wood Dale, IL) and 0.2mM Cholesteryl Hydrogen Succinate (MP Biomedicals) in 1X phosphate-buffered saline)) containing Streptavidin was injected into the chamber to coat the biotinylated surface with streptavidin molecules. After an incubation period, unbound streptavidin was washed away, and purified, fluorophore-labeled CLC-ec1 protein sample was applied to the slide. After the sample was allowed to adhere to the slide, the chamber was washed with reactive oxygen species (ROXS) scavenging solution (3.3% w/w glucose (SigmaAldrich), 3 units/mL pyranose oxidase (Sigma-Aldrich), 0.001% w/w catalase (Sigma-Aldrich), 1 mM ascorbic acid (Sigma-Aldrich), and 1 mM methyl viologen (Sigma-Aldrich), in Imaging Buffer (described above). The pH of the solution was adjusted with 0.1M Citric acid or 0.1M NaOH to desired pH value just before the slides were imaged. Details on replicates for each experimental condition are shown in **Table 2**.

**TABLE 2.** smFRET Replicate Summary

| <u>Protein</u> | <u>pH</u> | <u># of protein preparations</u> | <u># of labeling reactions</u> | <u># of slides</u> |
| --- | --- | --- | --- | --- |
| WT<sup>a,b</sup> | 7.5 | 3 | 9 | 11 |
|  | 6.5 | 1 | 2 | 4 |
|  | 5.5 | 1 | 3 | 6 |
|  | 4.5 | 3 | 8 | 12 |
|  | 4.0 | 2 | 3 | 6 |
| QQQ<sup>a</sup> | 7.5 | 1 | 6 | 9 |
|  | 4.5 | 1 | 6 | 9 |
| WT-QQQ<sup>a</sup> | 7.5 | 1 | 5 | 5 |
|  | 4.5 | 1 | 4 | 5 |
| QQQ-WT<sup>a</sup> | 7.5 | 1 | 3 | 6 |
|  | 4.5 | 1 | 3 | 6 |
| 75C/234C<sup>c</sup> | 7.5 | 3 | 7 | 10 |
|  | 4.5 | 3 | 6 | 9 |
| 75C/377C<sup>c</sup> | 7.5 | 2 | 6 | 12 |
|  | 4.5 | 2 | 6 | 12 |

<sup>a</sup>protein samples labeled at positions 377C/234C.
<sup>b</sup>in total, five independent WT 377C/234C protein preparations were examined.
<sup>c</sup>WT background

### smFRET Data Collection and Analysis

Data were collected using a PicoQuant Micro Time 200 confocal microscope. Excitation laser light from 532nm (LDH-D-TA-530; Picoquant, Berlin, Germany) and 637nm (LDH-D-C-640; Picoquant) lasers was used to excite the donor and acceptor fluorophores, respectively. A Pulsed Interleaved Excitation (PIE) setup was used with a pulse rate of 80 MHz to alternate the donor and acceptor excitation wavelengths. Excitation light traveled through an objective lens (100× 1.4 NA; Olympus, Tokyo, Japan) and interacted with the sample molecules. Emission light was collected through 550 nm (FF01-582/64;AHF, Tübingen-Pfrondorf, Germany /Semrock, Rochester, NY) and 650 nm (2XH690/70;AHF) emission filters and was then detected by two Single-Photon Avalanche Diodes (SPCM CD3516H; Excelitas technologies, Waltham, MA). Data were collected at a time resolution of 1 millisecond and then binned to 5 millisecond resolution to increase the signal-to-noise ratio. smFRET traces were obtained by measuring the fluorescence intensity of donor and acceptor fluorophores when excited at donor absorbance. Control traces were obtained for acceptor fluorescence with acceptor excitation.

Molecules that showed a single photobleaching step in each of the donor and acceptor channels and that also exhibited anticorrelation between the donor and acceptor fluorescence intensities were selected for analysis. The donor and acceptor intensity traces for selected molecules were subjected to corrections to eliminate background fluorescence, subtract signal resulting from bleed-through between the donor and acceptor channels, and account for differences in quantum yield or detector efficiency. The corrected donor (I_D_) and acceptor (I_A_) intensities were used to calculate the FRET efficiency (E_FRET_) using the following equation:

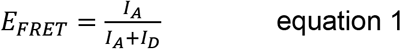

The combined corrected efficiency traces from all the molecules were used to evaluate the mean FRET efficiencies reported in **Table 3** and to generate the histograms of FRET efficiencies exhibited under different conditions. Gaussian curves were fit to the observed data using Origin software (OriginLab).

**TABLE 3.**
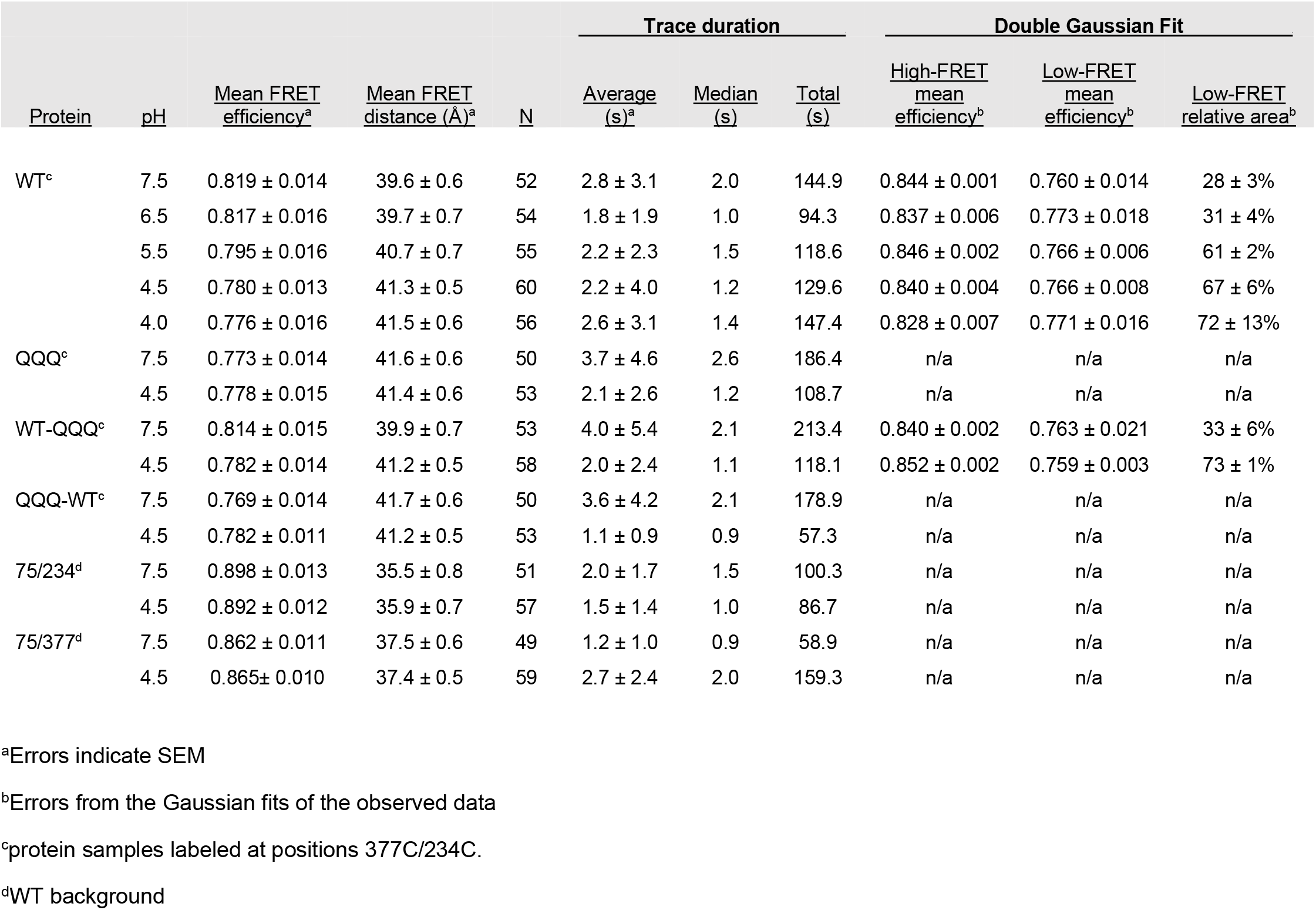
smFRET Summary Data

| Protein | pH | Mean FRET<br>efficiency<sup>a</sup> | Mean FRET<br>distance (Å)<sup>a</sup> | N | Trace duration |  |  | Double Gaussian Fit |  |  |
| --- | --- | --- | --- | --- | --- | --- | --- | --- | --- | --- |
|  |  |  |  |  | Average<br>(s)<sup>a</sup> | Median<br>(s) | Total<br>(s) | High-FRET<br>mean<br>efficiency<sup>b</sup> | Low-FRET<br>mean<br>efficiency<sup>b</sup> | Low-FRET<br>relative area<sup>b</sup> |
| WT<sup>c</sup> | 7.5 | 0.819 ± 0.014 | 39.6 ± 0.6 | 52 | 2.8 ± 3.1 | 2.0 | 144.9 | 0.844 ± 0.001 | 0.760 ± 0.014 | 28 ± 3% |
|  | 6.5 | 0.817 ± 0.016 | 39.7 ± 0.7 | 54 | 1.8 ± 1.9 | 1.0 | 94.3 | 0.837 ± 0.006 | 0.773 ± 0.018 | 31 ± 4% |
|  | 5.5 | 0.795 ± 0.016 | 40.7 ± 0.7 | 55 | 2.2 ± 2.3 | 1.5 | 118.6 | 0.846 ± 0.002 | 0.766 ± 0.006 | 61 ± 2% |
|  | 4.5 | 0.780 ± 0.013 | 41.3 ± 0.5 | 60 | 2.2 ± 4.0 | 1.2 | 129.6 | 0.840 ± 0.004 | 0.766 ± 0.008 | 67 ± 6% |
|  | 4.0 | 0.776 ± 0.016 | 41.5 ± 0.6 | 56 | 2.6 ± 3.1 | 1.4 | 147.4 | 0.828 ± 0.007 | 0.771 ± 0.016 | 72 ± 13% |
| QQQ<sup>c</sup> | 7.5 | 0.773 ± 0.014 | 41.6 ± 0.6 | 50 | 3.7 ± 4.6 | 2.6 | 186.4 | n/a | n/a | n/a |
|  | 4.5 | 0.778 ± 0.015 | 41.4 ± 0.6 | 53 | 2.1 ± 2.6 | 1.2 | 108.7 | n/a | n/a | n/a |
| WT-QQQ<sup>c</sup> | 7.5 | 0.814 ± 0.015 | 39.9 ± 0.7 | 53 | 4.0 ± 5.4 | 2.1 | 213.4 | 0.840 ± 0.002 | 0.763 ± 0.021 | 33 ± 6% |
|  | 4.5 | 0.782 ± 0.014 | 41.2 ± 0.5 | 58 | 2.0 ± 2.4 | 1.1 | 118.1 | 0.852 ± 0.002 | 0.759 ± 0.003 | 73 ± 1% |
| QQQ-WT<sup>c</sup> | 7.5 | 0.769 ± 0.014 | 41.7 ± 0.6 | 50 | 3.6 ± 4.2 | 2.1 | 178.9 | n/a | n/a | n/a |
|  | 4.5 | 0.782 ± 0.011 | 41.2 ± 0.5 | 53 | 1.1 ± 0.9 | 0.9 | 57.3 | n/a | n/a | n/a |
| 75/234<sup>d</sup> | 7.5 | 0.898 ± 0.013 | 35.5 ± 0.8 | 51 | 2.0 ± 1.7 | 1.5 | 100.3 | n/a | n/a | n/a |
|  | 4.5 | 0.892 ± 0.012 | 35.9 ± 0.7 | 57 | 1.5 ± 1.4 | 1.0 | 86.7 | n/a | n/a | n/a |
| 75/377<sup>d</sup> | 7.5 | 0.862 ± 0.011 | 37.5 ± 0.6 | 49 | 1.2 ± 1.0 | 0.9 | 58.9 | n/a | n/a | n/a |
|  | 4.5 | 0.865 ± 0.010 | 37.4 ± 0.5 | 59 | 2.7 ± 2.4 | 2.0 | 159.3 | n/a | n/a | n/a |

<sup>a</sup>Errors indicate SEM
<sup>b</sup>Errors from the Gaussian fits of the observed data
<sup>c</sup>protein samples labeled at positions 377C/234C.
<sup>d</sup>WT background

The distance between the two fluorophores was calculated with the following Förster equation:

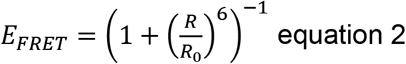

where R is the inter-dye distance, and R0 is the Förster radius, which we have determined to be 51 Å for the Alexa555-Alexa647 pair.

### Statistical Analysis

The mean (μ) FRET efficiencies were determined from the combined corrected FRET efficiency trace. Standard deviation (σ) was determined using eq 3,

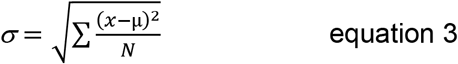

where N is the total number of data points in the combined FRET efficiency trace from all the molecules probed for a given condition. P values were calculated using mean, standard deviation, and number of molecules studied under a given condition, using unpaired t-tests.

### Summary of Supplemental Material

Supplemental materials include three figures showing additional examples of raw smFRET traces (Supplemental Figures 1 and 2) and a comparison of residues 233 and 234 on WT and QQQ CLC-ec1 (Supplemental Figure 3). Also included is a zip file containing csv files for all smFRET experiments described in this manuscript.

## RESULTS AND DISCUSSION

### Strategy for measuring smFRET within a single subunit

To investigate distances within a given subunit and to ensure no crosstalk between subunits, we co-expressed a cysteine-less construct containing a twin-strep tag (Schmidt et al., 2013) and a second construct with two cysteines and no strep tag. The purified protein sample containing the mixture of homodimeric cysteine-less/strep-tagged protein, homodimeric double-cysteine protein, and heterodimers was labeled with a mixture of thiol-reactive donor and acceptor fluorophores. This labeled mixture was *in situ* purified on a microscope slide coated with streptavidin, resulting in attachment of homodimeric cysteine-less protein and heterodimeric protein containing one cysteine-less subunit and one double-cysteine subunit (**Figure 2A**). The homodimeric cysteine-less protein has no fluorophores appended; hence, the fluorescence signal arises from only the heterodimeric protein. Heterodimeric protein labeled with donor-acceptor fluorophores can be isolated from protein with donor-donor and acceptor-acceptor labels in the smFRET measurement because proteins with donor-donor labels have no signal in the acceptor channel, and proteins with acceptor-acceptor labels have no signal in the acceptor channel (Dolino et al., 2017).

### Strategy to monitor the conformation of the extracellular gate

Helix N (**Figure 2B**) lines the anion-selectivity filter (Dutzler et al., 2002) and forms an extracellular bottleneck that controls Cl^−^ accessibility to and from the extracellular side of the protein (Khantwal et al., 2016). Its position changes in QQQ (outward-facing state) relative to WT (occluded state), producing an opening of the permeation pathway (**Figure 2B**). To monitor this movement, we placed fluorophore labels at position 377 on Helix N and at position 234 on Helix I, which is located on the opposite side of the permeation pathway from Helix N (**Figure 2C**), at a distance suitable for smFRET measurements, using the Alexa555 and Alexa 647 donor-acceptor pair (R~40 Å, given that R_0_ = 51 Å).

### smFRET monitors the status of the extracellular gate

We prepared two CLC-ec1 samples for smFRET measurements, one with the WT CLC-ec1 background and one with the QQQ mutant background, both fluorescently labeled on Helices N and I as shown in **Figure 2C**. These labeled samples (with double-cysteine mutations) will be referred to as “WT” and “QQQ” going forward. We measured smFRET of these samples at pH 7.5 and pH 4.5. The low-pH condition was used to favor protonation of the three key Glu residues (**Figure 1B**), so as to shift the conformational equilibrium of the WT protein towards the outward-facing state (Bell et al., 2006; Elvington et al., 2009; Abraham et al., 2015; Khantwal et al.; Chavan et al., 2020). Representative traces showing the fluorescence intensity from the donor and acceptor channels, together with the calculated FRET efficiencies, are shown in **Figure 3A** and **Supplemental Figure 1**. Since the CLC-ec1 transporter turnover rate (~2000 s^−1^) (Walden et al., 2007) is much faster than our smFRET sampling rate (200 s^−1^), the smFRET traces represent a weighted average of any conformational states present. Intriguingly, we did observe a few smFRET traces in which the FRET efficiency was seen to switch between states; however, these events were too rare for methodical analysis (**Supplemental Figure 2**).

The cumulative histograms from 50-60 molecules for each sample condition are shown in **Figure 3B**. For the WT protein, the cumulative histograms show that there is a shift in the population to lower FRET efficiency when the pH is reduced from pH 7.5 to pH 4.5. This low-FRET histogram for WT at pH 4.5 resembles the smFRET histograms for the QQQ protein at both pH conditions. These results indicate that the smFRET monitors the WT protein population shifting towards a QQQ-like (outward-facing) conformation at low pH. From the FRET efficiencies, the mean distance between the probes corresponds to 39.8 and 41.3 Å for the high- and low-FRET states respectively (**Table 3**). Although this distance change is in the opposite direction expected based on the C-alpha carbon distances in the WT and QQQ protein structures (39.0 Å and 38.1 respectively), this discrepancy is not surprising given that the FRET measurement reflects distances between tethered fluorophores (not C-alpha carbons) together with the assumptions involved in converting FRET values to absolute distances (Lerner et al., 2020). The important point is that the FRET measurement follows the predicted change: lowering the pH shifts the WT FRET signal, and this shift is in the direction of the FRET signal observed with the QQQ protein, thus providing confidence that a shift of the population towards the outward-facing conformation is being monitored. In addition, we note that the distance change is in the predicted direction if we evaluate distances between helices N and I instead of C-alpha positions of the labeled residues (**Supplemental Figure 3**).

**Figure 2.**
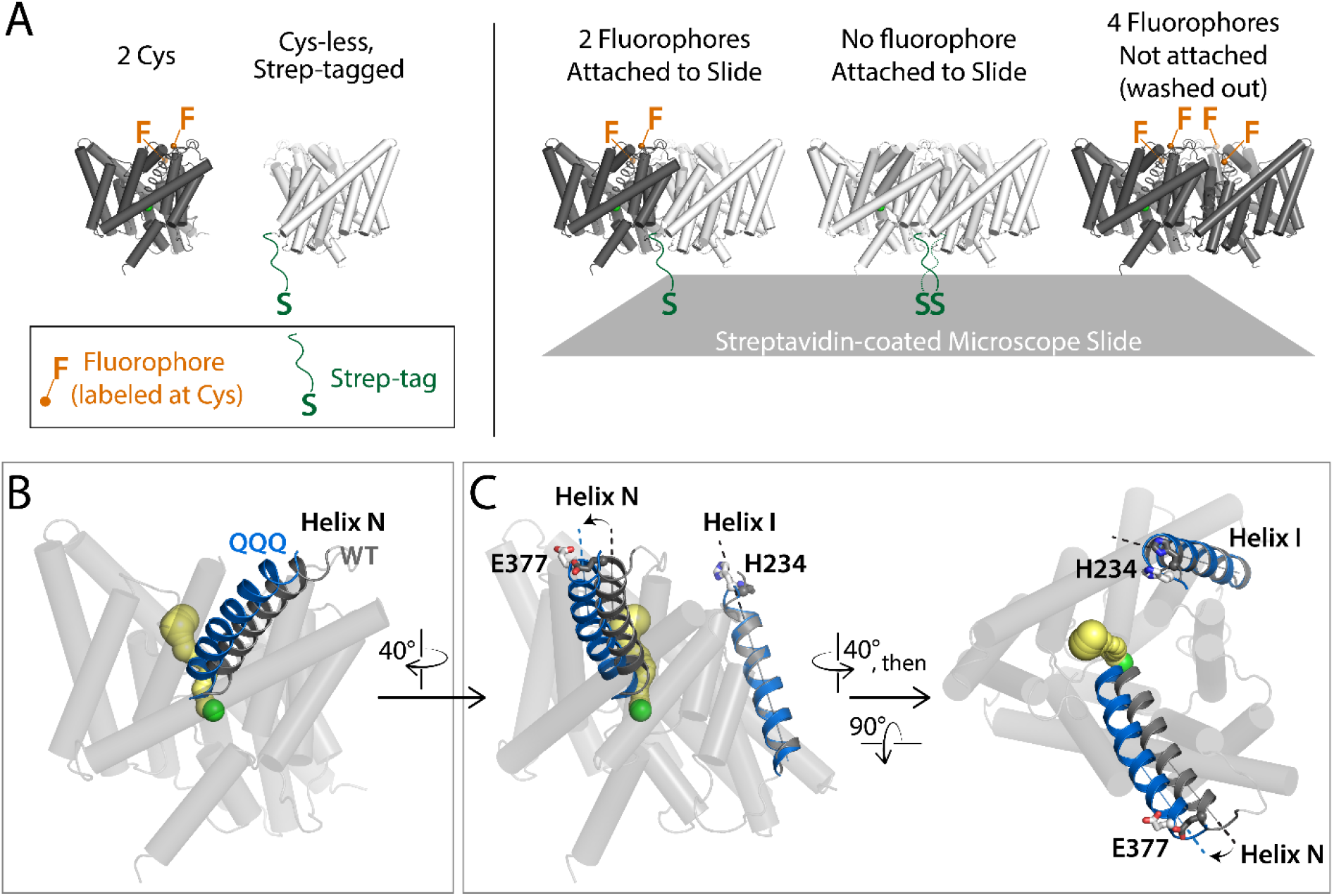
Experimental Strategy to monitor the CLC-ec1 external vestibule. (**A**) *In situ* purification to obtain samples with fluorophores on only one subunit. Two CLC-ec1 constructs (double-cysteine mutant and Strep-tagged cysteine-less) were co-expressed, purified, and labeled with a mixture of thiol-reactive donor (Alexa 555) and acceptor (Alexa 647) fluorophores, both depicted in orange. To obtain samples with only one subunit labeled, a streptavidin-coated microscope slide was used to pull down strep-tagged proteins. (**B**) Helix N, shown in ribbon, changes position in the outward-facing QQQ structure (blue) compared to the occluded WT structure (grey). This movement contributes to the opening of the permeation pathway in the QQQ structure (yellow Caver pathway, see also **Figure 1B**). (**C**) Labeling strategy to monitor the status of the extracellular gate. Fluorophores covalently modifying cysteines at E377C (Helix N) and H234C (Helix I) allow smFRET measurements to monitor the Helix-N gate-opening movement.

**Figure 3.**
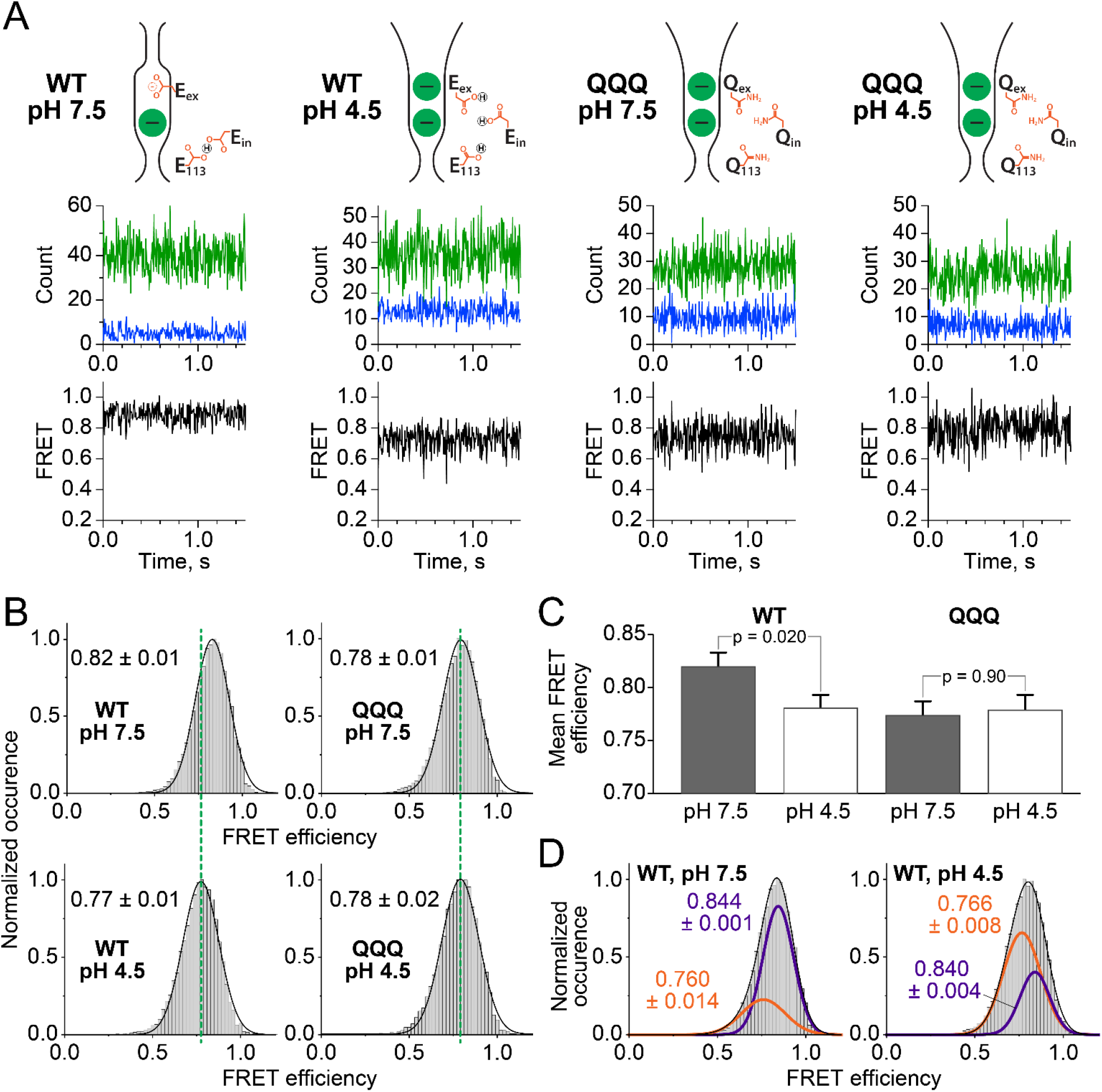
smFRET of CLC-ec1 labeled at 377C/234C. (**A**) smFRET data were obtained for constructs in either the WT or QQQ background, at pH 7.5 and pH 4.5. The cartoons at the top of each panel indicate the preferred conformation of the extracellular gate and key glutamate residues (**Figure 1B**) under each condition. For each condition, a representative raw data trace is shown; green is acceptor intensity, blue is donor intensity, black is FRET. Additional representative traces are shown in Supplemental Figure 1, and the full data set for each set of experiments are available as supplemental material. (**B**) Histograms for each sample show the normalized occurrence of FRET efficiency values. The mean FRET efficiency (average ± SEM) for each condition is indicated next to each histogram. (**C**) Bar graph plot of the mean FRET efficiency. In the WT background, there is a significant shift with pH, to the FRET efficiency level observed with QQQ at both pH conditions. (**D**) Fits of the WT smFRET data to double Gaussian functions reveal distinct high-FRET and low-FRET distributions, fits shown in purple and orange, respectively (mean and error from the fitting algorithm). At pH 7.5, the fits yield mean FRET efficiencies of 0.84 and 0.76, with the low-FRET distribution containing 28% of the population (**Table 3**). At pH 4.5, the mean FRET values for the two Gaussians are the same as observed at pH 7.5, but the distributions have shifted, with the low-FRET distribution containing 67% of the population (**Table 3**).

These results demonstrate that smFRET can monitor conformational change involving small (~1- Å) motions. Typically, smFRET is used to monitor conformational changes involving distance changes in the 10 – 20 Å range (Lerner et al., 2018)), which is compatible with the scale of conformational change in most transporters (Zhao et al., 2010; Forrest et al., 2011; Akyuz et al., 2015; Dyla et al., 2017; Han et al., 2017; Ren et al., 2018; de Boer et al., 2019; Zhu et al., 2019). In contrast, the conformational change from occluded to outward-facing state in CLC-ec1 involves changes on the order of 1.0 – 2.5 Å (Chavan et al., 2020). Within the constraints of the smFRET experiment (selecting labeling positions separated by ~40 Å and avoiding positions where mutation is known to alter function), the larger distances within this range were not accessible and hence we targeted the 377/234 (Helix N/I) pair. Inspired by our success with this pair (**Figure 3**), we tested additional labeling pairs around the extracellular permeation pathway, on Helices C-N and C-I. For the Helix C-N (75/377) labeling pair, the structures predict no significant change between occluded and outward-facing states (**Figure 4A**), and the smFRET data are consistent with this prediction (**Figure 4B**). For the Helix C-I (75/234) labeling prediction, the structures predict a small change, of similar magnitude to the change predicted for the Helix N-I (377/234) pair (**Figure 4A**). However, this predicted change was not in our smFRET assay (**Figure 4B**). Though disappointing, this result is in keeping with the fact that stochastic movements of the FRET probes (Antonik et al., 2006), and other issues already mentioned regarding the relationship between FRET efficiency and absolute distance, can make it difficult detect small (~1-2 Angstrom) conformational changes. Nonetheless, we are fortunate in having identified a pair of labelling positions (377/234) that *does* report on conformational changes at this scale, as confirmed by the QQQ mutant. Our success in monitoring conformational change using one of the two pairs with predicted changes provides a practical proof of concept that smFRET can be used to monitor the conformational state of a transporter than undergoes relatively small movements.

**Figure 4.**
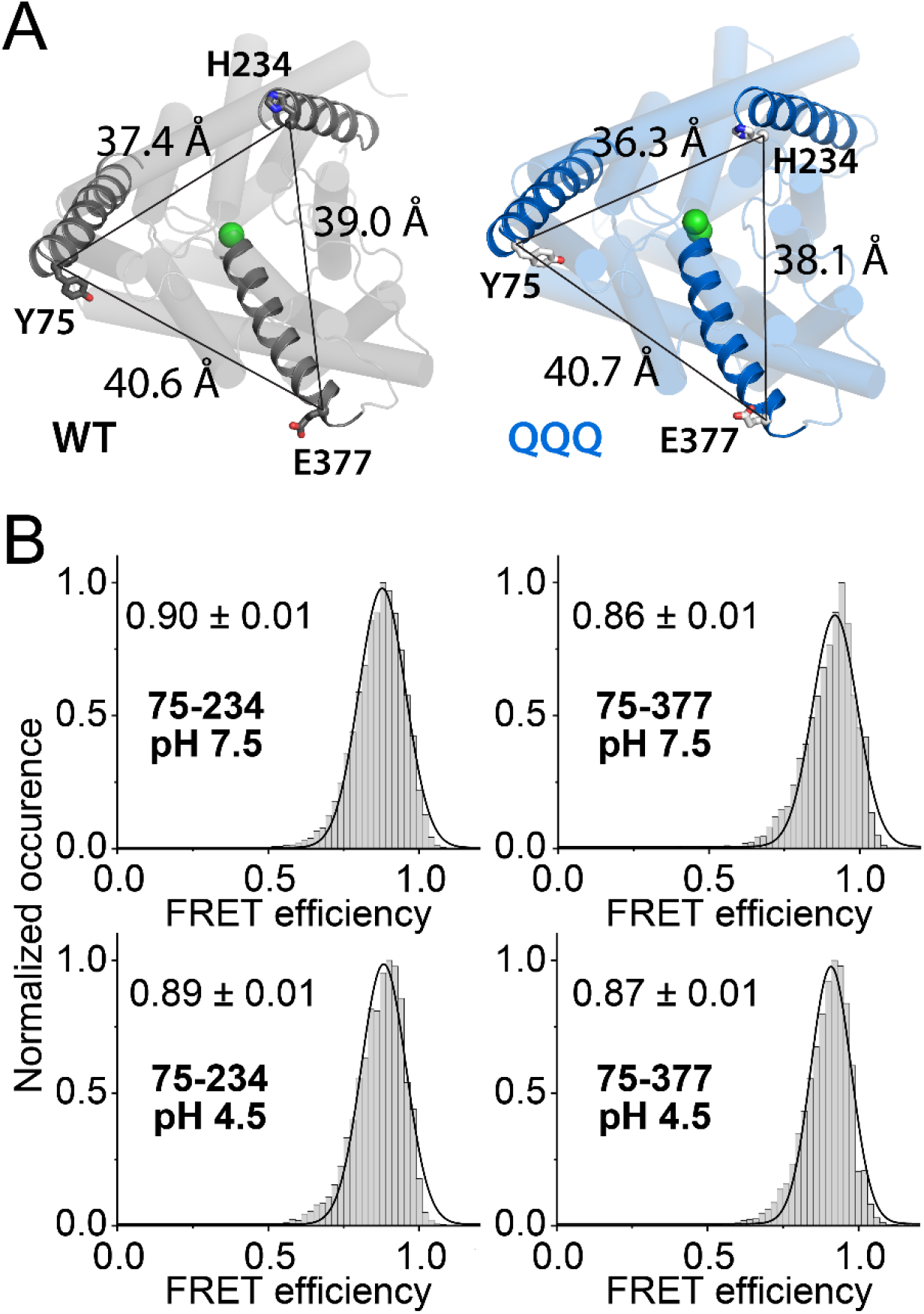
smFRET of CLC-ec1 labeled at positions 75C/234C and 75C/377C. (**A**) Diagram of the triangulation of labeling positions designed to measure CLC-ec1 conformational change at the extracellular side. (**B**) smFRET summary data for the 75C/234C and 75C/377C labeling pairs. See also **Table 3**. Unlike the 377C/234C, FRET measurements with these pairs showed no significant change with pH.

### Relationship to the CLC transport cycle

The smFRET data for 377/234 labeling on both WT and QQQ samples are well fit to single Gaussian functions, consistent with the existence of a single population of conformational states under each condition (pH 7.5 or pH 4.5). However, given the statistically significant difference in FRET efficiency between the probes in the two conditions (**Figure 3C**), and the fact that CLC transport is rapid, we hypothesized that the data might reflect the average of a rapidly fluctuating mixed population, and therefore we also evaluated fits of each data set to a double Gaussian function. Here, the WT and QQQ data differ markedly. For QQQ at either pH condition, fitting the data to a double Gaussian fit yields a second Gaussian that is not significantly different in FRET efficiency from the first. In contrast, for WT the double Gaussian fits yield distinct distributions with mean FRET efficiencies of 0.84 and 0.76 (**Figure 3D, Table 3**). Further, the relative population shifts with pH, with the low-FRET population increasing from 28 ± 3% at pH 7.5 to 67 ± 6% at pH 4.5 (**Figure 3D, Table 3**). These data suggest that the WT protein populates both occluded and outward-facing states under both conditions, with the outward-facing state becoming favored at low pH.

Secondary active transport requires an “alternating access” mechanism, in which a transporter’s binding sites for substrates are alternately accessible to the extracellular and intracellular solutions (Jardetzky, 1966; Tanford, 1983; Forrest et al., 2011). The CLC transport cycle is uniquely complex (Miller and Nguitragool, 2009), as CLCs are the only secondary active transporters known to exchange an anion for a cation. However, the basic requirement for inward- and outward-facing conformational states remains (Basilio et al., 2014). A simplified model of the CLC Cl^−^/H^+^ exchange cycle is shown in **Figure 5A**. Cl^−^/H^+^ exchange is achieved as follows. Starting from the top left and moving clockwise, Cl^−^ from the intracellular side enters the Cl^−^ permeation pathway, displacing Glu_ex_ (“E_ex_”) via a knock-off mechanism (Miller and Nguitragool, 2009). Subsequent protonation of Glu_ex_ favors a conformational change to the outward-facing state, which opens the Cl^−^ pathway to the outside and the H^+^ pathway to the inside (Chavan et al., 2020). Following release of H^+^ to the intracellular side, the deprotonated Glu_ex_ returns to the Cl^−^ pathway, knocking Cl^−^ out to the extracellular side. This transport cycle nets exchange of 2 Cl^−^ for 1 H^+^, in either direction. Detailed variants of this model can be found in recent publications (Chavan et al., 2020; Leisle et al., 2020).

**Figure 5.**
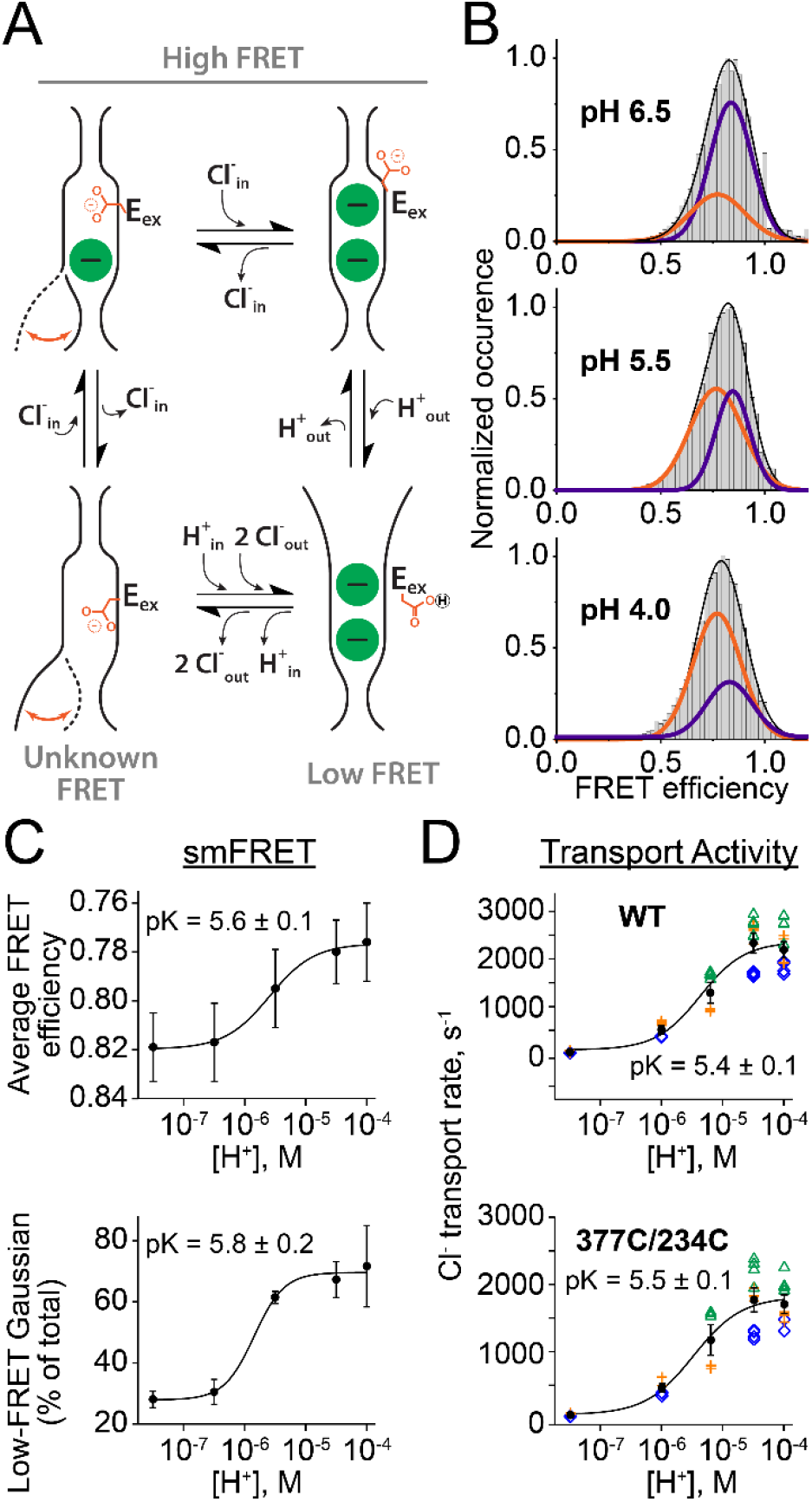
pH dependence of conformational equilibria and transport activity. (**A**) Simplified cartoon depiction of the CLC Cl^−^/H^+^ transport cycle, involving occluded (high-FRET), outward-facing (low-FRET), and inward-facing (unknown FRET) conformational states. Occluded states (top cartoons) are observed in most CLC-ec1 crystal structures, including WT CLC-ec1, where Glu_ex_ is in the Cl^−^-permation pathway (*top left cartoon*) and E148Q CLC-ec1, where the Gln_ex_ (mimicking protonated Glu_ex_) is rotated upwards (top right cartoon). The outward-facing state is observed in the QQQ crystal structure (Chavan et al., 2020) (bottom right cartoon). For the inward-facing state (bottom left cartoon), there is currently no high-resolution structure; however, the involvement of a conformational change at the inner gate is supported by cross-linking (Basilio et al., 2014) and spectroscopic (Bell et al., 2006; Abraham et al., 2015) studies. (**B**) smFRET efficiency histograms for the labeled WT protein at pH 4.0, 5.5, and 6.5, showing fits to double Gaussian functions, as in **Figure 3C**. (**C**) smFRET data (*top*, average FRET value; *bottom*, relative area of low-FRET Gaussian) are plotted as a function of [H^+^]. The solid lines are fits of the data to the equation: Y = Y_max_/(1+(K_a_/[H^+^])^n^), where Y_max_ is the maximum for the y-axis signal, K_a_ is the apparent affinity for H^+^, and n is the Hill coefficient. For the average-FRET data, n = 1.2 ± 0.4 and pK_a_ = 5.6 ± 0.1; for the relative-area data, n=1.8 ± 0.7 and pK_a_ = 5.8 ± 0.2. (**D**) pH dependence of transport activity for WT (*upper graph*) and 377C/234C (*lower graph*). Data are from experiments performed on 3 independent protein preparations, as indicated by the different colors. Black symbols indicate the average transport rate at each [H^+^], ± SEM. Data are fit as in panel C. For WT, n = 1.2 ± 0.4 and pK_a_ = 5.4 ± 0.1; for 377C/234C, n = 1.1 ± 0.3 and pK_a_ = 5.5 ± 0.1.

The transport-cycle model highlights an important limitation to interpreting our smFRET data. Since the molecular structure of the inward-facing conformational state is unknown, we do not know whether or how this state contributes to the smFRET signal. Keeping this caveat in mind, we investigated the pH dependence of the smFRET change in more detail. smFRET histograms acquired for the WT labeled sample at pH conditions ranging from 4.0 to 7.5 show that the peak of the smFRET efficiency shifts from 0.819 to 0.776 as the pH is decreased from 7.5 to 4.0 (**Figure 5B, Table 3**). When fit to double Gaussian functions, the relative area of the low-FRET peak increases from 28 to 71% as the pH is decreased from 7.5 to 4.0 (**Figure 5C, Table 3**). These changes (mean smFRET or relative area from double-Gaussian function) can be fit to a Hill equation with a pK_a_ value of 5.6 and 5.8 respectively (**Figure 5C**). Notably, this pH dependence closely parallels the pH dependence of transport activity (**Figure 5D**). While it would be imprudent at this stage to impose a mechanistic interpretation on this correlation (for the reason mentioned above as well as the fact pH influences both conformational equilibria and substrate concentration), its existence suggests that the smFRET measurement is monitoring a conformational change relevant to the transport cycle. Consistent with this idea, the lack of effect of pH on QQQ conformation (**Figure 3A,B**) correlates with the lack of effect of pH on QQQ transport rates (Chavan et al., 2020). We anticipate that future studies, once labels have been developed to monitor the inward-facing state, will provide deeper insight into the relationship between CLC-ec1 conformational equilibria and the transport mechanism.

### Independence of the subunits

We applied the smFRET method to evaluate the independence of the CLC-ec1 subunits. In CLC channels and transporters, ion permeation and gating occur independently within each subunit of the homodimer (Accardi, 2015; Jentsch and Pusch, 2018). In addition to this independent functioning, many CLC homologs – both channels and transporters – exhibit a cooperative gating mechanism that regulates the two subunits simultaneously (Miller, 1982, 2014; Jentsch and Pusch, 2018), and mutations in one subunit can affect gating conformational changes of a wild-type subunit in the adjacent subunit (Lorenz et al., 1996; Pusch, 2002). For CLC-ec1, the ability of subunits to act as independent transporters has been confirmed by the characterization of a monomerized variant (Robertson et al., 2010). However, whether transport in CLC-ec1 dimers is regulated by a cooperative gating mechanism remains unknown. To evaluate this possibility, we used our *in situ* labeling strategy (**Figure 2A**) to generate 2 sets of heterodimers, one with fluorophore-labeled QQQ and an adjacent unlabeled WT subunit, and a second with fluorophore-labeled WT and an adjacent unlabeled QQQ subunit. smFRET was measured at pH 7.5 and pH 4.5 for both samples. For the WT-QQQ sample (WT labeled, QQQ unlabeled), the smFRET data show a shift with pH that can be fit with a double Gaussian (**Figure 6A**). This shift is not measurably different from that observed with the WT homomeric sample (**Figure 6C, Table 3**). Similarly for the QQQ-WT sample (QQQ labeled, WT unlabeled), the smFRET signal is not influenced by the adjacent subunit: the data are well fit by a single Gaussian that is not influenced by the shift from pH 7.5 to 4.5 (**Figure 6B,C, Table 3**). Thus, there is no measurable cooperativity in the conformational transition monitored in our smFRET assay.

**Figure 6.**
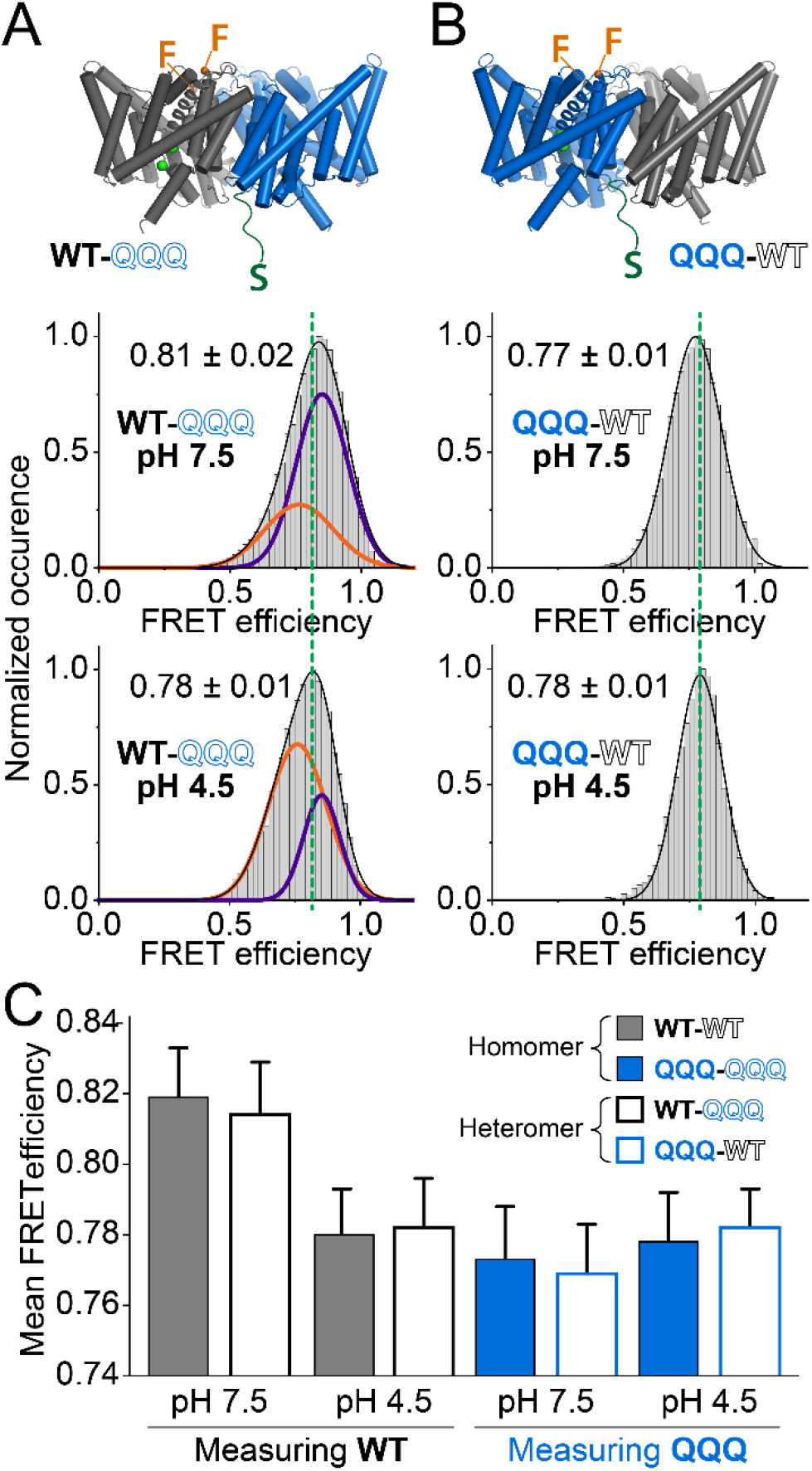
smFRET state is not influenced by the adjacent subunit. smFRET cumulative histograms for (**A**) labeled WT subunit with adjacent unlabeled QQQ and (**B**) labeled QQQ subunit with adjacent unlabeled WT, at pH 7.5 (*top panels*) and pH 4.5 (*bottom panels*). (**C**) Summary bar graph showing that the labeled WT and QQQ mean FRET distance (± SEM, see **Table 3**) is unaffected by the nature of the adjacent subunit. Mean FRET.

### Summary/Conclusion

We have developed a single-molecule FRET approach for monitoring conformational change in a CLC-ec1 subunit without crosstalk from the second subunit. We applied this approach to monitor CLC-ec1 conformational change in response to changes in [H^+^]. Taking advantage of the CLC-ec1 QQQ mutant, which adopts an outward-facing conformation, we confirmed that the conformational transition from occluded to outward-facing states is being monitored. By using heteromeric constructs with WT and QQQ background, we showed that this conformational transition occurs independent of the adjacent subunit. Going forward, smFRET measurements using additional labeling positions and following conformational change under different conditions (varying Cl^−^, using well-characterized mutants (Walden et al., 2007; Jayaram et al., 2008; Lim and Miller, 2009), and using samples reconstituted into liposomes), will be a powerful approach for understanding the relationship between CLCs’ conformational dynamics and transport mechanisms.

## ACKNOWLEDGMENTS

We thank Martin Prieto for comments on the manuscript. This research was funded by the National Institutes of Health GM113195 (M.M.), GM122528 (V. J.), and F31GM130035 (R.J.D.).

The authors declare no competing financial interests.

## AUTHOR CONTRIBUTIONS

Ricky Cheng, Ayush Krishnamoorti, Vladimir Berka, and Ryan Durham performed experiments. Ricky Cheng, Ayush Krishnamoorti, Vasanthi Jayaraman, and Merritt Maduke wrote the manuscript. Merritt Maduke and Vasanthi Jayaraman supervised research. All authors contributed to data analysis, experimental design and interpretation, and editing/revising the manuscript.

---

**Supplemental Figure 1.**
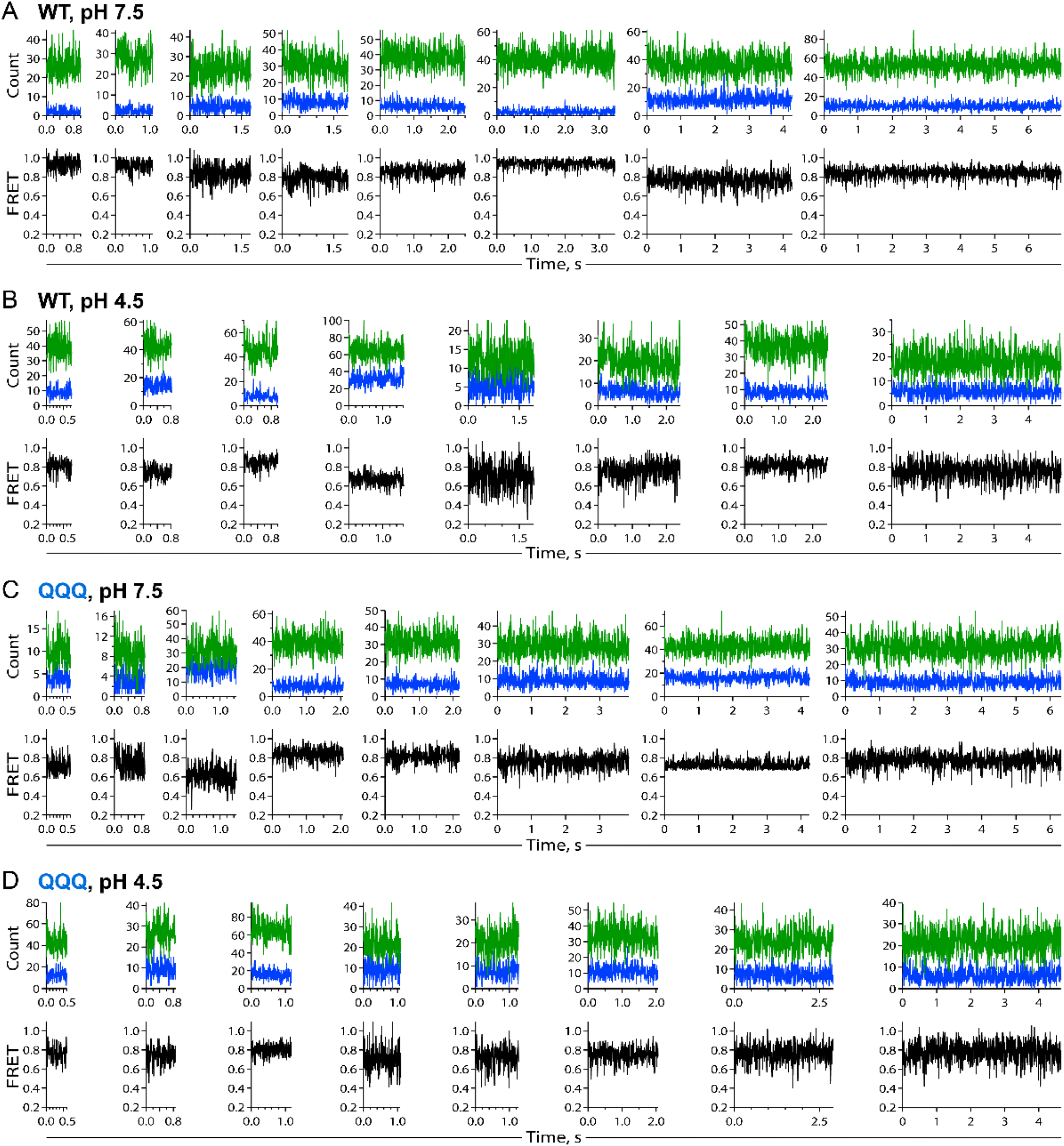
Additional representative smFRET traces for the samples labeled at 377C/234C. (**A**) WT pH 7.5. (**B**) WT pH 4.5 (**C**) QQQ pH 7.5 (**D**) QQQ pH 4.5

**Supplemental Figure 2.**
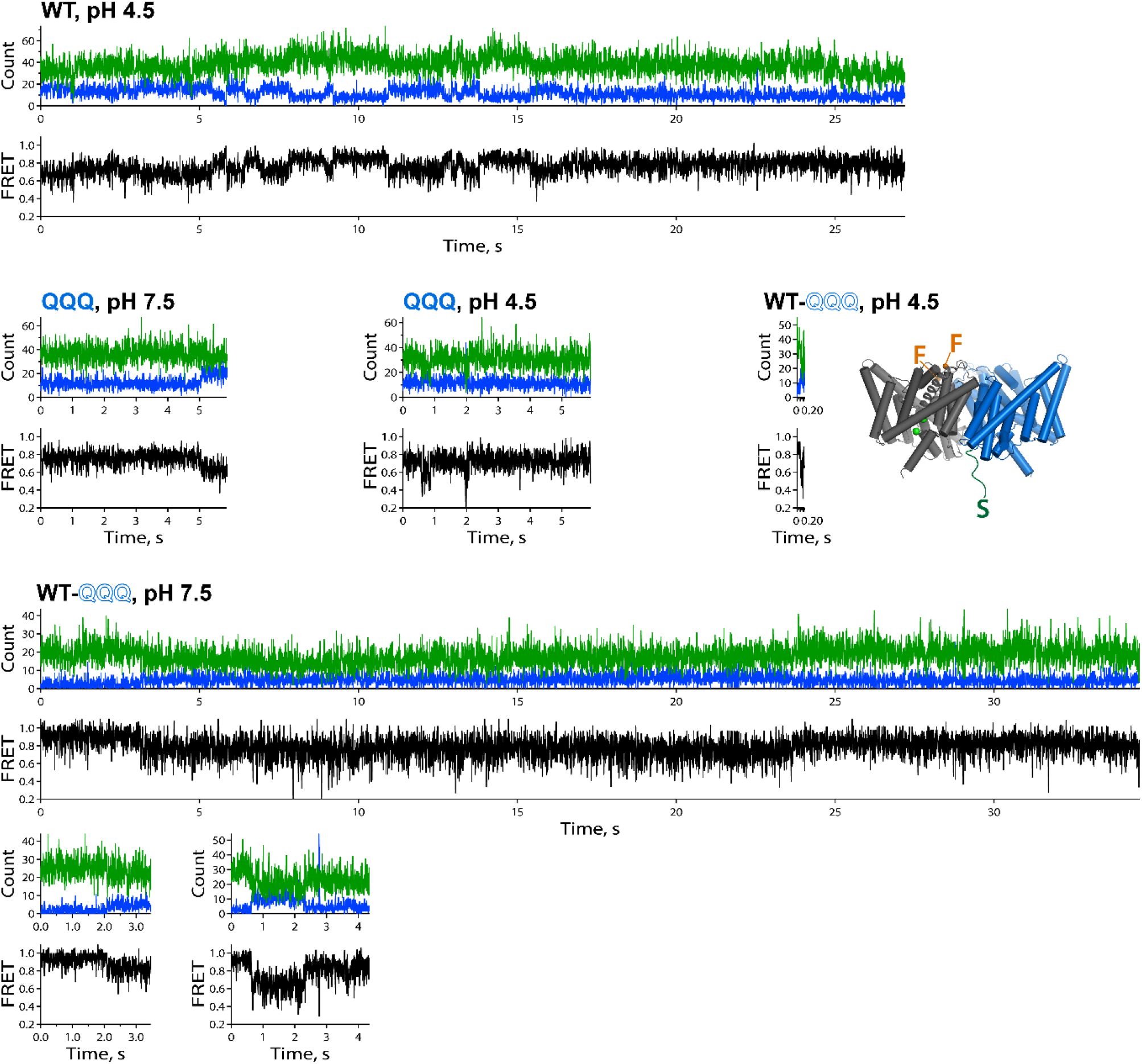
smFRET traces showing transitions between FRET-efficiency levels. Sample/conditions are indicated.

**Supplemental Figure 3.**
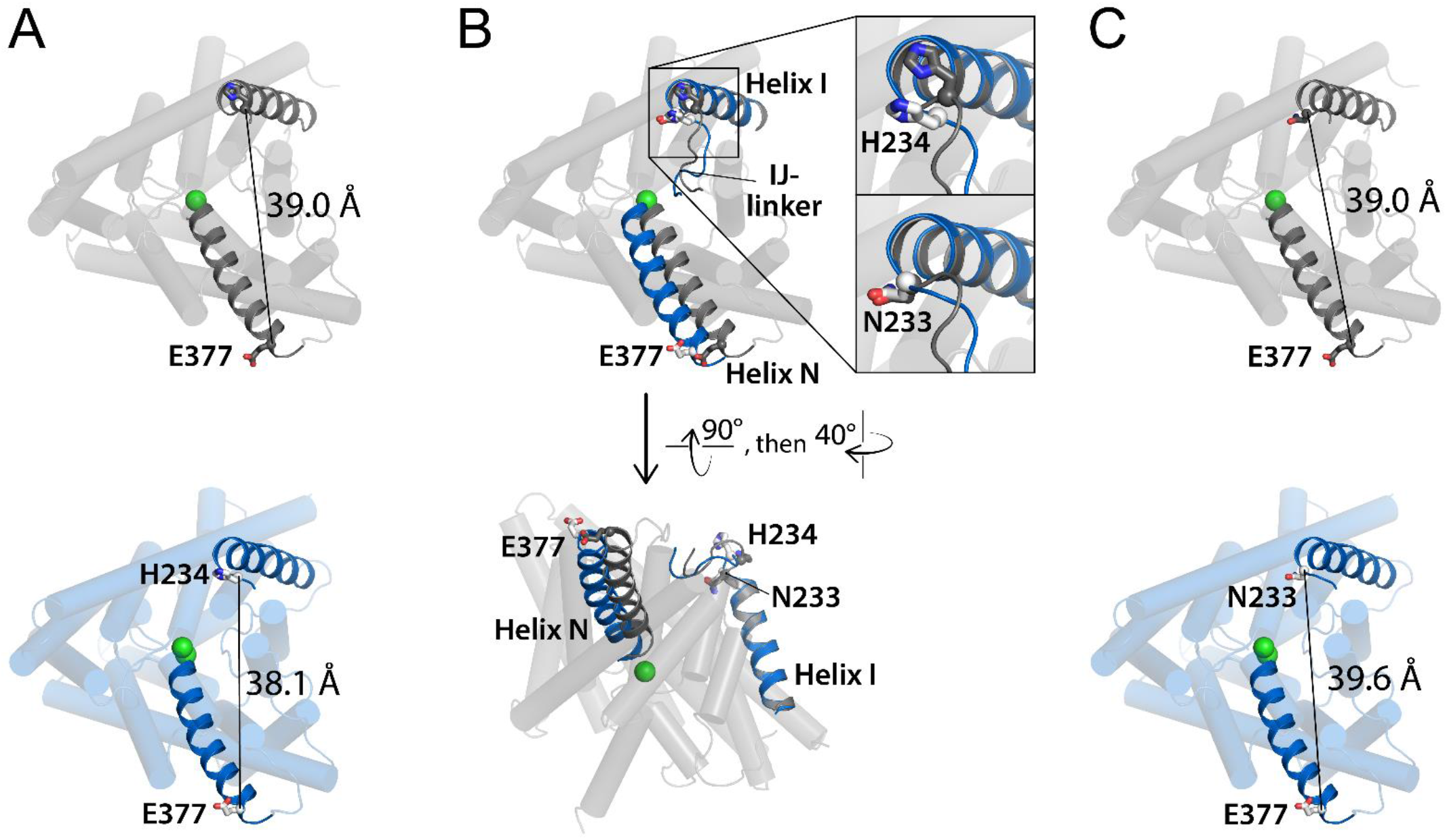
Helix N moves away from Helix C. (**A**) The distance between the Cα atoms on residue 377 (last residue on Helix N) and residue 234 (first residue *after* Helix I) decreases by 0.9 Å in QQQ compared to WT CLC-ec1. Protein subunits are shown viewed from the extracellular side. (**B**) Side view showing that the distance between Helix N and Helix I *increases* in QQQ compared to WT CLC-ec1. (**C**) The distance between the Cα atoms on residue 377 and residue 233 (the first residue on Helix I) increases by 0.6 Å in QQQ compared to WT CLC-ec1. This distance change is of similar magnitude and direction to the distance change reported by our smFRET labels on residues 377C/234C (**Table 3**). The mutant N233C is inactive, and therefore we used H234C instead for the FRET labeling experiments.

## Notes

### Competing Interest Statement

The authors have declared no competing interest.

